# Phenotypic consequences of RNA polymerase dysregulation in *Escherichia coli*

**DOI:** 10.1101/150581

**Authors:** Paramita Sarkar, Amy Switzer, Christine Peters, Joe Pogliano, Sivaramesh Wigneshweraraj

## Abstract

Many bacterial adaptive responses to changes in growth conditions due to biotic and abiotic factors involve reprogramming of gene expression at the transcription level. The bacterial RNA polymerase (RNAP), which catalyzes transcription, can thus be considered as the major mediator of cellular adaptive strategies. But how do bacteria respond if a stress factor directly compromises the activity of the RNAP? We used a phage-derived small protein to specifically perturb bacterial RNAP activity in exponentially growing *Escherichia coli*. Using cytological profiling, tracking RNAP behavior at single-molecule level and transcriptome analysis, we reveal that adaptation to conditions that directly perturb bacterial RNAP performance can result in a biphasic growth behavior and thereby confer the ‘adapted’ bacterial cells an enhanced ability to tolerate diverse antibacterial stresses. The results imply that while synthetic transcriptional rewiring may confer bacteria with the intended desirable properties, such approaches may also collaterally allow them to acquire undesirable traits.

## INTRODUCTION

Bacteria use a variety of adaptive strategies that allow them to survive and persist in the face of unfavorable growth conditions. Global alterations in gene expression underpin many such strategies. These are often mediated and controlled at the transcriptional level through the modulation of the activity and specificity of the transcriptional machinery, the RNA polymerase (RNAP). Therefore, the plasticity of the bacterial adaptive transcriptional response to unfavorable growth conditions is often mediated by the action of *cis* and *trans* acting regulatory factors that modulate RNAP activity and the associations between the RNAP and the different promoter-specificity sigma (σ) factors (reviewed in (1)). In *Escherichia coli*, the transcription of genes during the exponential growth phase is carried out by the RNAP containing the σ^70^ factor (Eσ^70^), while Eσ^38^ (one of the six alternative σ factors) is required to execute the global adaptive transcriptional changes in response to diverse stress signals that impede growth, including the transition from exponential to stationary phase of growth. Since any adaptive response to changes in growth conditions begins with a reprogramming of cellular activity at the transcriptional level, the bacterial RNAP can be considered as the major mediator of bacterial adaptive responses. The phenotypic consequences of transient perturbation to the transcription programme through factors that directly compromise the activity and specificity of the RNAP (such as action of bacteriocins, bacteriophage (phage)-encoded proteins, aberrant transcription factor activity or synthetic rewiring of the transcriptional programme) are sparsely understood. Gp2 is a 7 kDa T7 phage protein, which binds to the *E. coli* RNAP tightly and efficiently inhibits Eσ^70^ activity *in vitro* (reviewed in (2)). However, unlike rifamycin, the antibiotic which inhibits RNA chain elongation by the RNAP indiscriminately of its σ factor composition, Gp2 can be regarded as a more selective inhibitor of the bacterial RNAP: While Eσ^70^ is efficiently inhibited by Gp2, RNAP containing alternative σ-factors, σ^38^ or σ^54^, are relatively less sensitive to this inhibition – although, the efficacy of Eσ^38^ is compromised by Gp2 *in vitro* (3,4). This is also consistent with the principal biological role of Gp2 during T7 infection, which is to prevent aberrant Eσ^70^ activity on the phage genome during phage replication and packaging of phage virions (5). However, since the *E. coli* chromosome becomes degraded by T7-encoded endonucleases shortly after the inhibition of the host RNAP by Gp2, any effect of Gp2 on the host transcriptome during successful T7 infection is unlikely to be physiologically relevant to the phage. Therefore, we used Gp2 as a molecular tool to synthetically compromise RNAP activity and thereby induce perturbations to the transcriptional programme in exponentially growing *E. coli* to study how bacteria adapt and respond to such conditions.

## MATERIALS AND METHODS

### Growth Assay

Details of the bacterial strains and plasmids used in the study are listed in Table S1. Bacteria were grown in Luria-Bertani (LB) broth medium at 37 °C, 180 rpm. Growth assays on multi-well platforms were conducted in 48 well plates (Greiner) using 500 μl culture volume and quantified using a BMG Labtech SPRECTOstar Nano microplate reader. Growth assays in larger batch cultures were conducted using 100 ml culture volume and the growth profile was obtained by hourly measurement of optical density (OD_600nm_). The seed cultures generated for all the growth assays were supplemented with 0.2% (v/v) glucose along with the appropriate antibiotic (ampicillin at 100 μg/ml and chloramphenicol at 35 μg/ml) and 1:100 dilution of the seed culture was used to inoculate the bacterial cultures used for determining the growth profiles. Gp2 overexpression was induced (with either 0.2% (v/v) L-arabinose or 1 mM IPTG) ~2 h after inoculation at OD_600nm_ values ranging from 0.2-0.4 (depending the strain used) when the cells were in the exponential phase of growth.

### Pull-down assays

*E. coli* MC1061 transformed with pBAD:Gp2 was grown as described above and the time points at which samples (50 ml) were taken are indicated in the figures. The MagneHis^TM^ Protein Purification System (Promega) was used for pull-down assays according to manufacturer’s instructions. Briefly, the bacterial pellet was resuspended in 3 ml of Tris Buffered Saline (TBS) and lysed by sonication using a Sonics Vibracell sonicator (settings: amplitude 40%, 5 sec on, 5 sec off pulse for 5 min). The lysate was then mixed with 100 μl of resin beads and incubated for 1 h at 4°C. The beads were washed with 5x resin bed volume of TBS. To elute bound proteins from the beads, 100μl of 2x SDS sample buffer (0.125 M Tris-HCl, pH 6.8, 4% (w/v) SDS, 20% (v/v) glycerol, 10% (v/v), 2-mercaptoethanol, 0.004% (w/v) bromophenol blue) was added and the beads were boiled for 1 min at 100 °C. Ten microliters of the eluted sample was loaded on a 4-20% gradient SDS-polyacrylamide electrophoresis (PAGE) gel and proteins of interest were detected by Western blotting (see below).

### Western Blotting

Samples from the pull-down assays were separated by SDS-PAGE and transferred to Polyvinyl difluoride (PVDF) membrane (0.45 μm) using Trans-Blot^®^ Turbo^TM^ Transfer System device and the blots were processed according to standard Western blotting protocols. For detection of proteins in lysates, total protein was precipitated using Trichloroacetic acid (TCA). The TCA pellet was treated with 0.2 mM NaOH for 10 min at room temperature and dissolved in 200 μl solubilizing buffer (8 M Urea, 0.1 mM Dithiothreitol). The protein extract was mixed with 2x SDS sample buffer and 10 μl was used for SDS-PAGE and Western blotting. The titres of the primary antibodies used were as follows: anti-*E. coli* RNAP β-subunit antibody at 1:1000 [8RB13 - Abcam], anti- *E. coli* RNAP α-subunit antibody at 1:1000 [4RA2 – BioLegend] and anti-6X His tag^®^ antibody (HRP) at 1:5000 [ab1187 – Abcam], anti-*E. coli* RNAP β’-subunit antibody at 1:1000 [NT73 – BioLegend], anti- *E. coli* RNAP σ^70^-subunit antibody at 1:1000 [2G10 – BioLegend], anti- *E. coli* RNAP σ^38^-subunit antibody at 1:1000 [1RS1 – BioLegend], anti-DnaK antibody at 1:5000 [ab69617 – Abcam]. Rabbit Anti-Mouse IgG H&L (HRP) was used at 1:2500 [ab97046 – Abcam] (where necessary) as the secondary antibody. The blots were developed using the Amersham ECL Western Blotting Detection Reagent and analysed on a ChemiDoc. Digital images of the blots were obtained using an LAS-3000 Fuji Imager, and signal quantification and calculations were performed exactly as described by Shadrin et al. (6). Briefly, to estimate protein concentrations from the Western blot signals, a calibration curve was generated using known concentrations of α and Gp2-His6 using their respective antibodies. The amount of proteins at each time point was then estimated from the calibration curve.

### Bacterial Cytological Profiling

*E. coli* containing pBAD and pBAD:Gp2 were grown in LB medium as described above. Overnight cultures were diluted 1:100 into fresh LB and were grown at 37°C until the OD_600nm_ was measured to be ~0.2 at which point 0.2% (v/v) L-arabinose was added to induce overexpression of Gp2. Images were taken every hour for ten hours. Cells were stained with FM 4-64 (2 μg/ml) to visualise the membranes, DAPI (2 μg/ml) to visualise the DNA, and SYTOX Green (2 μg/mL), a vital stain, which is normally excluded from cells with an intact membrane but brightly stains cells that are lysed (7). Cells were visualised on an Applied Precision DV Elite optical sectioning microscope equipped with a Photometrics Cool- SNAP-HQ^2^ camera. Pictures were analysed using SoftWoRx v5.5.1 (Applied Precision). For DAPI quantification (Supplementary Figure S2), images were analysed using CellProfiler v. 2.1.1 to generate automated measurements of mean DAPI Intensity. Cells were initially identified using the FM 4-64 images, and the objects were expanded using the phase image as a guide to obtain final cell objects. The mean DAPI intensity for each cell object was measured. DAPI intensity of individual cells was quantitated (background DAPI intensity level was subtracted), binned into groups based on intensity, and the percentage of cells in each intensity bin was plotted.

### Photoactivated localization microscopy (PALM) and single particle tracking

For PALM and single particle tracking experiments an *E. coli* strain KF26 containing an endogenous fusion of PAmCherry to the β’ subunit of RNAP (8) was used, pBAD:Gp2 was transformed into this strain. The bacterial cultures were grown as described above and cells were sampled at the time points indicated in the text and imaged and analysed in a similar way as previously described (8,9). Briefly, 1 ml of cells were centrifuged, washed and resuspended in low fluorescence minimal media, 1 μl of this suspension was placed on a minimal medium agarose pad (10) and cells were imaged on a PALM-optimised Nanoimager (Oxford Nanoimaging, <u>www.oxfordni.com)</u> with 15 millisecond exposure over 10,000 frames for four separate experiments. Photoactivatable molecules were activated with a 405 nm laser and then imaged and bleached with a 561 nm laser. For RNAP mobility analysis, the Nanoimager software suite was first used to localise the activated molecules by finding intensity peaks that were significantly above background, then fitting the detected spots with a Gaussian function. The Nanoimager single-particle tracking feature was then used to map trajectories for the individual RNAP molecules over multiple frames, using a maximum step distance between frames of 0.3 μm and a nearest-neighbour exclusion radius of 1.2 μm. This feature reports apparent diffusion coefficient (D*) for the specified acquisition, using point-to-point distances in the trajectories.

### RNA sequencing

Cultures were grown as described above and sampled (50 ml) at the time points indicated in the figures and text. Two biological replicates were performed for each sample. Cell pellets were outsourced to Vertis Biotechnologie AG for further processing. Briefly, total RNA was isolated from the cell pellets using a bead mill and *mir*Vana^TM^ RNA isolation kit (Ambion) according to the manufacturer’s instructions. Ribosomal RNA (rRNA) from the total RNA sample was depleted using Ribo-Zero rRNA removal kit for bacteria (Epicentre). The rRNA-depleted RNA samples were fragmented using RNase III. Samples were then poly (A)-tailed followed by treatment with Tobacco Acid Pyrophosphatase (TAP, Epicentre). First strand cDNA synthesis was performed using oligo (dT)-adapter primer and M-MLV reverse transcriptase. Resulting cDNAs were PCR amplified. The primers used for PCR amplification were designed for TruSeq sequencing according to the manufacturer’s guidelines (Illumina). The cDNA was sequenced on the Illumina NextSeq 500 system. The data analyses were performed with the ‘CLC Genomics Workbench 7’ using standard parameters. RNA-seq reads were mapped to the *E. coli* K-12 MG1655 (U00096) genome using Burrows-Wheeler Aligner. Reads that mapped uniquely were used for further analysis. The number of reads mapping to each gene was calculated and matrix of read counts was generated. The matrix was analysed using the DESeq2 BioConductor package for differential gene expression analysis. Genes with ≤ 10 reads mapped to them were excluded from analysis. All statistical analyses were performed in R studio version 0.00.442.

### Antibacterial stress assays

For antibiotic sensitivity assays, 5 ml of bacterial culture was collected at specified time points during growth in 50 ml Corning conical centrifuge tubes. The culture was then treated with gentamicin (50 μg/ml, 10X MIC for *E. coli*) or ciprofloxacin (10 μg/ml, 10X MIC for *E. coli*) and the culture was allowed to grow at 37°C for 5 hours. Two hundred microliters samples were collected at various times (indicated in Figure 5) following antibiotic treatment, washed and serially diluted in TBS. To quantify colony-forming units (CFU) individual dilutions were plated on LB agar plates and CFUs were counted. To calculate log10 percentage survival the ratio of CFU of untreated cells to and treated cells was obtained and multiplied by 100 for each time point. The H_2_O_2_, low pH and osmotic stress assays were done as previously described (11-13). Briefly, for H_2_O_2_ challenge, samples were collected as above but resuspended in LB containing 42 mM H_2_O_2_ and incubated for 1 hour at 37°C shaking at 180 rpm. The reaction was stopped with 2μg/ml catalase and processed as above. For pH challenge, samples were collected as above, resuspended in LB with pH 54.5 and the culture was incubated for 1hr at 37°C shaking at 180 rpm and processed as above. For osmotic shock challenge, samples were collected as above, resuspended in LB containing 0.6 M NaCl and the culture was incubated for 2 hours at 42°C shaking at 180 rpm and processed as above. To calculate % survival the ratio of CFU of untreated cells and treated cells at a given time point was multiplied by 100.

## RESULTS AND DISCUSSION

### Overexpression of recombinant Gp2 in exponentially growing *E. coli* induces a biphasic growth pattern

Using a multi-well platform, we compared the growth properties of *E. coli* strain MC1061 (which is able to transport arabinose, but is unable to metabolize it; (14)) in which Gp2 overexpression was induced with L-arabinose from an *araBAD* promoter (pBAD:Gp2) with that of *E. coli* cells exposed to different concentrations of rifamycin. As shown in Figure 1A, the induction of Gp2 expression in exponentially growing *E. coli* cells (phase A) resulted in the rapid attenuation of growth. Strikingly, following a 6-7 hour period of stasis (phase B), the cells recovered growth (phase C), albeit at a much slower rate of growth than phase A cells. In marked contrast, the recovery phase was not observed in *E. coli* cells exposed to different concentrations of rifamycin under identical growth conditions. We repeated the experiment in 100 ml batch cultures and plotted the growth curve as Log_10_OD_600nm_ against time (h) to accurately measure the rate of growth in phase A (before Gp2 induction) and in phase C (during recovery); here from on all growth curves are shown in this format. As shown in Figure 1B, the rate of growth in phase C was ~10-fold less than that in phase A. Identical results were obtained when we repeated the experiment with *E. coli* strain MC1061 in which expression of Gp2 was under the control of an IPTG inducible *T5* promoter or in a pBAD plasmid (pBAD33) in which the ampicillin-resistance conferring gene was replaced with the gene conferring chloramphenicol-resistance, suggesting that recovery of growth is not due to depletion of L-arabinose or an indirect effect of ampicillin, respectively (Supplementary Figure S1A). Since *E. coli* strain MC1061 contains functionally deleterious *relA* and *spoT* genes (15), which control the synthesis and degradation of the major bacterial stress alarmone guanosine penta/tetraphosphate ((p)ppGpp), we investigated whether the growth behaviour by *E. coli* strain MC1061 in response to Gp2 induction is due to its inability to synthesis sufficient (p)ppGpp. Therefore, we compared the growth characteristics of *E. coli* strain MC1061 with that of *E. coli* strain BW25113 (which contains functional *relA* and *spoT* genes; (16)) in response to Gp2 overexpression. Results shown in Supplementary Figure S1B clearly show that *E. coli* strain BW25113 responds identically to Gp2 overexpression as the *E. coli* strain MC1061. Analysis of whole-cell extracts prepared from *E. coli* strain MC1061 phases B and C cells by Western blotting revealed that Gp2 was present at ~4-fold excess over the RNAP (Figure 1C), suggesting that the depletion of Gp2 did not lead to recovery of growth. Consistent with this observation, the addition of L-arabinose to phase C cells did not lead to the attenuation of growth as seen with phase A cells (Supplementary Figure S1C). We next considered whether the *E. coli* cells in phase C could have acquired genetic changes that have made them refractory to Gp2. Results from two independent experiments suggest that this is unlikely to be the case: In the first experiment, to test if either the plasmid-borne recombinant Gp2 or plasmid pBAD:Gp2 itself has acquired any functionally-deleterious mutations that rendered Gp2 inactive or prevented Gp2 from being expressed, respectively, we isolated pBAD:Gp2 from phase C cells, transformed it into fresh *E. coli* MC1061 cells and grew the freshly transformed cells to exponentially phase; upon induction with L-arabinose, we detected the expected growth arrest as previously seen (Supplementary Figure S1D). In the second experiment, we considered the possibility that phase C cells represent an enriched sub-population of cells that were genetically resistant to inhibition by Gp2. To exclude this possibility, we harvested phase C cells, extensively washed them with phosphate-buffered saline (to remove any carry-over L-arabinose) and used them to re-inoculate fresh growth media. We found that the recovered cells responded to induction with L-arabinose in an identical manner as fresh cells (Supplementary Figure S1E), suggesting that observed recovery of growth is not due to the appearance of a subpopulation of Gp2-resistant cells (also see below).

**Figure 1.**
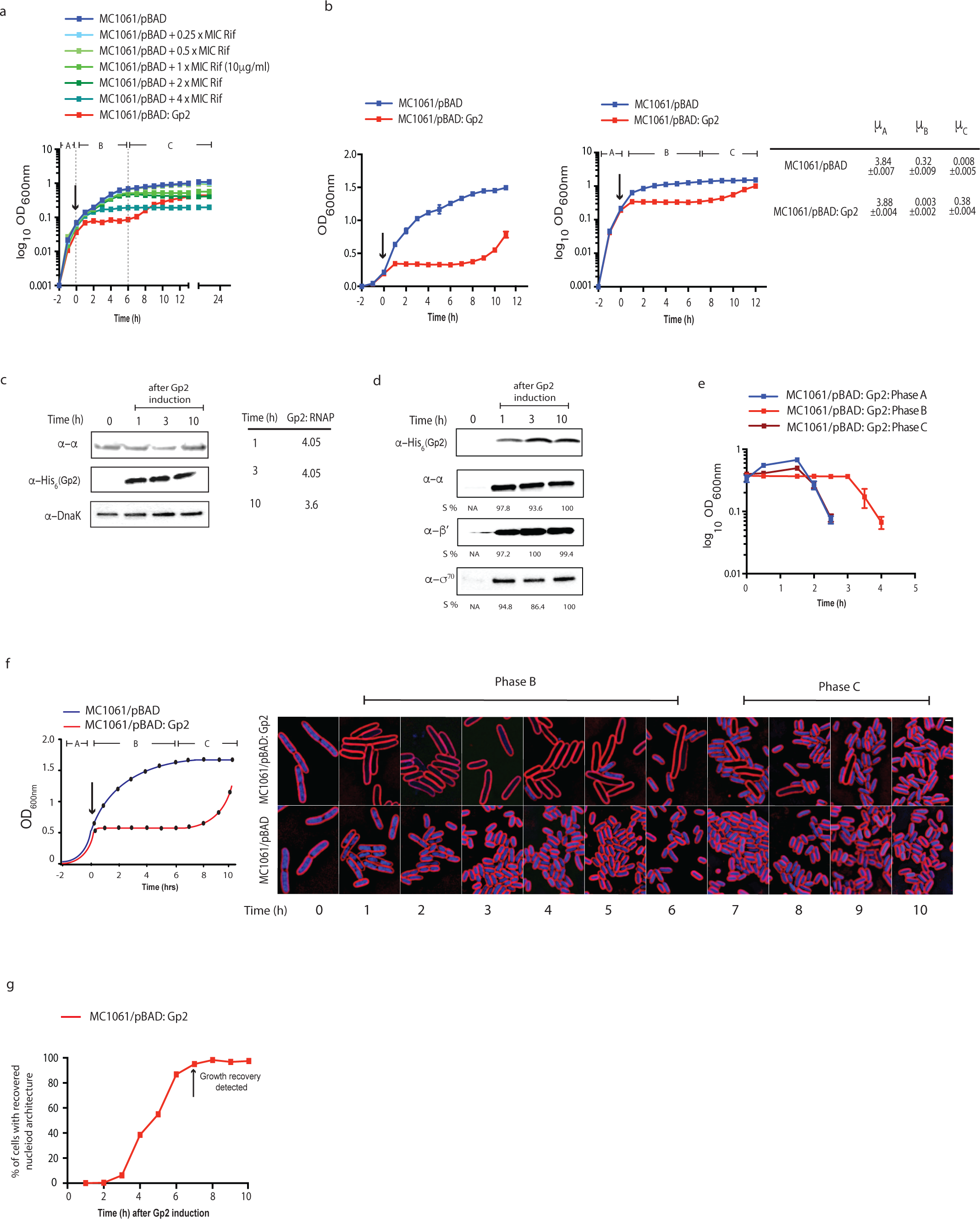
Overexpression of recombinant Gp2 in exponentially growing *E. coli* induces a biphasic growth behaviour. **(A)** Graph showing (log_10_ of the optical density (OD_600nm_) as function of time (hour)) of a culture of *E. coli* MC1061 cells containing pBAD or pBAD:Gp2 exposed to various concentrations of rifamycin (Rif) (MC1061/pBAD cells only) or L-arabinose (MC1061/pBAD:Gp2 cells only). The arrow indicates when either rifamycin or L-arabinose was added to the culture. **(B)** As in (A), but graphs show OD_600nm_ value (left) and log_10_ of the OD_600nm_ value (right) as function of time. On the graph on the right, the three distinct phases of growth seen with cells overexpressing Gp2 are indicated and the growth rates (μ) for each phase (A-C) are given next to the graph. **(C)** Image of a Western blot showing the relative abundance of Gp2 and RNAP (here the a-subunit is used to as a surrogate for RNAP) in whole cell extracts of cells from the indicated time points following induction of Gp2 overexpression; DnaK serves as a loading control. The table on the right of the Western blot image shows the relative ratio of Gp2 to RNAP at the indicated time point. **(D)** Image of a Western blot showing that the relative amount of Gp2 bound Eσ^70^ in remains unchanged at the indicated time points following induction of Gp2 overexpression (see text for details). The %S values for a, β’ and σ^70^ bands at each time point is calculated relative to the signal corresponding to the Gp2 bands from the same time point and the maximum signal intensity for a, β’ and σ^70^ is taken as 100%. **(E)** Graph showing OD_600nm_ value as a function of time of phase A, phase B and phase C cells following infection with T7 phage (see text for details). **(F)** Fluorescence microscopy images of cytological profiles of *E. coli* cells containing MC1061/pBAD and MC1061/pBAD:Gp2 before and after induction with Larabinose. The time points at which the cells were harvested for analysis is shown in the schematic growth curve on the left of the microscopy images. (**G**) Graph showing the percentage of cells with cytological profiles which indicate recovery of transcription activity (see text for details) as a function of time after induction of Gp2 overexpression. The percentage of recovered cells at each time point was calculated by quantifying DAPI intensity of individual cells, cells were binned into groups based on intensity and each intensity bin is plotted as a function of time. The arrow indicates the time point when growth recovery was detected.

To investigate whether intracellular conditions in phase C cells were unfavourable for Gp2 to effectively interact with (and therefore inhibit) the Eσ^70^ (which is presumably the RNAP form driving growth recovery) we compared the amount of Eσ^70^ that co-purifies with Gp2 in phase C with the amount of Eσ^70^ that co-purifies with Gp2 in phase B by Western blotting the samples using an antibody against the α, β’ and σ^70^ subunits of the RNAP. Results show no discernible differences in the amount of α, β’ and σ^70^ subunits *ispo facto* Eσ^70^ that co-purified with Gp2 from both phase B and C cells, suggesting that Gp2 is able to interact equally well with the RNAP in the stasis and recovery phases (Figure 1D). Further, since Gp2 activity is essential for T7 development and the eventual lysis of *E. coli* cells (5), we compared the ability of wild-type T7 phage to infect and lyse phase C cells to indirectly demonstrate that the intracellular condition in phase C is not unfavourable for Gp2 to bind to and inhibit the RNAP. As shown in Figure 1E, cell from phases A and C were effectively lysed by wild-type T7 phage 90 minutes following infection under our conditions.

To further understand the physiological changes occurring over time with Gp2 overexpression, we examined individual cells using Bacterial Cytological Profiling (BCP). BCP relies upon quantitative fluorescence microscopy to observe changes in cells exposed to antibiotics. Antibiotics that inhibit different pathways generate different cytological responses that can be interpreted using a database of control antibiotics (7). We used BCP to analyse individual cells in the three phases (A-C). Within 1 hour of Gp2 induction with L-arabinose, cells appeared similar to rifamycin treated cells, as expected since they both inhibit RNAP (Figure 1F and Figure S2A). Cells were slightly elongated and the nucleoids were decondensed, with dim DAPI fluorescence uniformly filling the cell (Figure 1F and Supplementary Figure S2B). As shown in Figure 1F, we detected recovery of nucleoid architecture and DAPI fluorescence within individual cells beginning at 3 hours, when 6% (n=192) of cells appeared similar to wild type. Recovery continued slowly, with 38% (n=340) recovered at 4 hours, ultimately reaching 97% at 10 hours post induction during recovery. Thus, it seems that cellular transcription activity resumes 3-4 hours after Gp2 overexpression, but visible growth recovery (judged by increase in OD_600nm_ value) only happens at 8-10 hours (Figure 1F). Overall, we conclude that Gp2 mediated dysregulation of RNAP activity results in a bi-phasic growth pattern, which, we suggest, is due to an adaptive response that occurs early in phase B that leads to the gradual resumption of growth in phase C in a heterogeneous manner.

### Overexpression of recombinant Gp2 in exponentially growing *E. coli* induces changes in RNAP behavior at single molecule level and induces global alterations in the transcriptional programme

We next focused on RNAP behavior and activity in response to Gp2 overexpression. Using photoactivated localization microscopy combined with single-particle tracking of individual RNAP molecules in live *E. coli* cells, we calculated the apparent diffusion coefficient (D*) from the mean squared displacement of trajectories of individual RNAP molecules in phases A-C. Since RNAP molecules engaged in transcription will have an overall slower diffusion rate because they are bound to DNA for longer periods of time than non-transcribing RNAP molecules that interact transiently with the DNA, we reasoned that changes to the D* value could be indicative of Gp2-induced changes in RNAP behavior in phases B and C. As shown in Figure 2A, the RNAP molecules in phase A cells had a D* value of 0.193 μm^2^s^-1^ (±0.010), whereas the RNAP molecules in phase B and C had higher D* values of 0.252 - 0.278 μm^2^s^-1^ (±0.032); this increased D* value was not seen in the presence of a functionally defective variant of Gp2-R56E (17) (Figure 2B), suggesting that the changes in the D* value we detected are due to the specific action of Gp2 on the RNAP. Overall, it seems that the overexpression of Gp2 clearly alters the behavior of transcribing RNAP molecules. Further since non-transcribing, thus non DNA-bound, RNAP molecules in *E. coli* have been reported to have D* value of 2-3 μm^2^s^-1^ under similar experimental conditions (8), the results also suggest that Gp2 does not cause the full dissociation of RNAP from the DNA.

**Figure 2.**
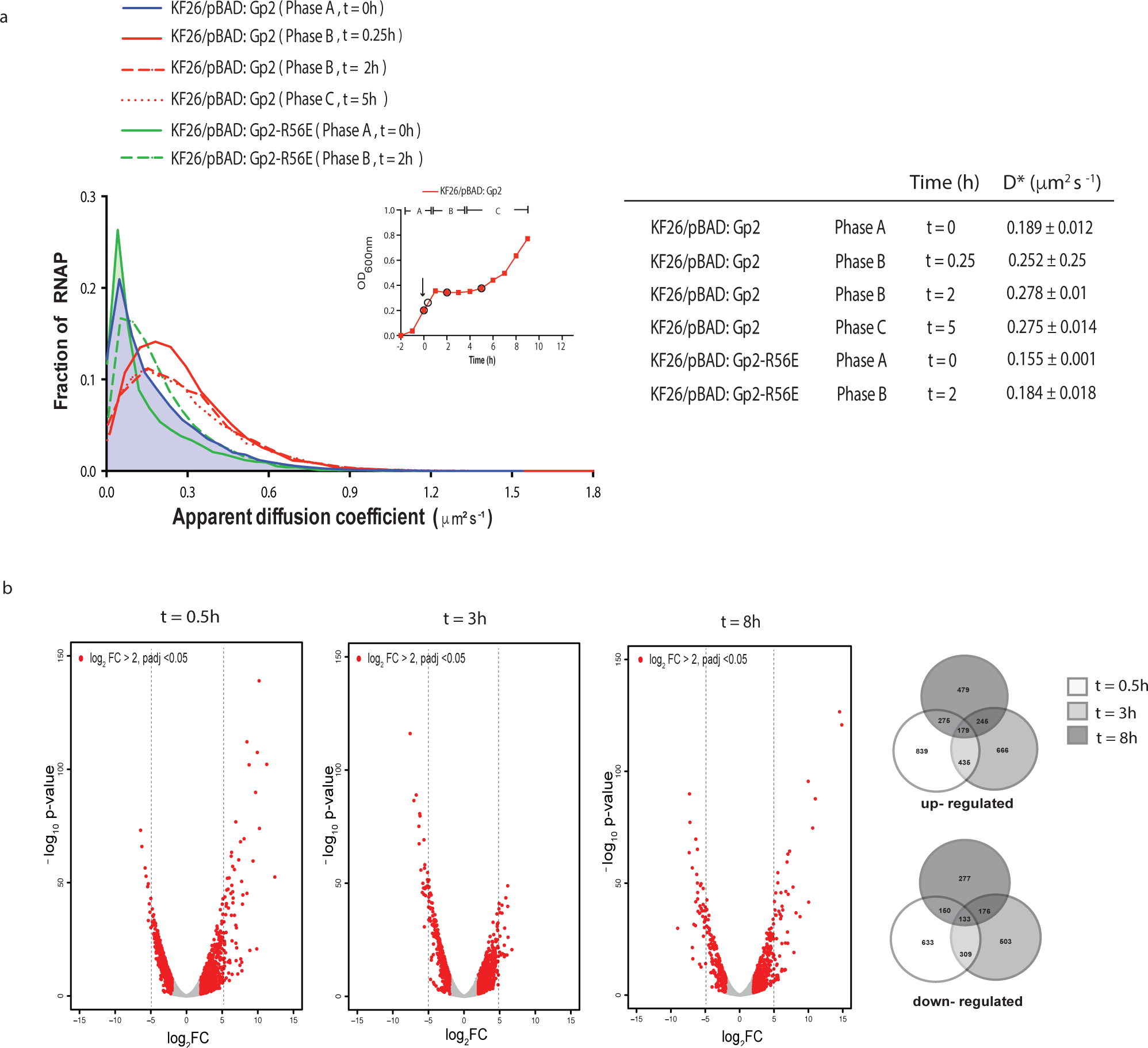
Overexpression of recombinant Gp2 in exponentially growing *E. coli* induces changes in RNAP behavior at single molecule level and induces global alterations in the transcriptional programme. **(A)** Graph showing the distribution of apparent diffusion co-efficient (D*) of RNAP molecules in *E. coli* KF26 cells containing pBAD, pBAD:Gp2 or pBAD:Gp2-R56E from phases A and B; The apparent diffusion co-efficient of each strain at the different phases is tabulated on the right (see text for details). The inset shows a graph of OD_600nm_ as a function of time of *E. coli* KF26 cells containing pBAD:Gp2 and the arrow indicates the time point when L-arabinose was added to the culture; the time points at which samples were obtained for analysis are circled. **(B)** The volcano plots show distribution of all differentially expressed genes with log2 fold change > 2 at t=0.5, t=3 (phase B) and t=8 (phase C) relative to gene expression at t=0. The number of genes up- and down-regulated at the different time points is represented in the Venn diagram on the right.

Next, to determine how the Gp2-induced changes in RNAP behavior affects the transcriptional programme of *E. coli*, we compared the global transcriptomes of phase B cells (at t=0.5 h and t=3 h post Gp2 induction) and phase C cells (t=8 h post Gp2 induction) with that of phase A cells obtained immediately prior to the addition of L-arabinose (t=0). We defined differentially expressed genes as those with expression levels changed ≥2-fold at t=0.5, 3 and 8 hours relative to t=0, with a False-Discovery-Rate-adjusted *p*-value <0.05. The volcano plots of gene expression generated following these criteria, clearly indicated that Gp2, induces significant alterations in the transcriptional programme of *E. coli* (Figure 2B): In phase B cells, at t=0.5, shortly after Gp2 expression and onset of stasis, a total of 1372 genes were differentially expressed of which, 839 and 633 were up- and down-regulated, respectively. Similarly, after 3 hours of stasis, a total of 1169 genes were differentially expressed, of which, 666 and 503 were up and down-regulated, respectively. Interestingly, consistent with the onset of growth recovery, in phase C, more genes were up-regulated (479) than down-regulated (277) with a total of 756 genes differentially expressed. Overall, the results clearly indicate that the overexpression of Gp2 in exponentially growing *E. coli* cells, an inhibitor of Eσ^70^, does not result in the full shut-off of transcription, but substantially alters RNAP behavior (Figure 2A) and thus the transcriptional programme (Figure 2B) of the cell.

### Adaptation to the Gp2 mediated perturbation to the transcriptional programme involves the action of several small non-coding regulatory RNAs

Bacterial adaptive strategies to stress can involve small non-coding regulatory RNAs that play important roles in the post-transcriptional regulation of gene expression and in the immediate response to stress and/or the recovery from stress. Strikingly, we note that 42 small RNA genes were some of the most highly up-regulated genes shortly after Gp2 overexpression (Figure 3A and Supplementary Figure S3). At t=0.5, 42 small RNA genes were up-regulated by more than five-fold compared to t=0 (Figure 3A and Supplementary Figure S3). Although most of small RNA genes were down-regulated at t=3 compared to t=0, we found a subset of them to be again up-regulated during recovery at t=8. Therefore, we considered whether small RNAs could contribute to how *E. coli* responds and adapts to Gp2 mediated perturbation to the transcriptional programme. Since a large number of these small regulatory RNAs depend on Hfq for pairing to their target mRNAs (indicated by red asterisks in Supplementary Figure S3), we compared the growth characteristics of *E. coli* strain BW25113:Δ*hfq* with that of the parent strain in response to overexpression of Gp2. As shown in Figure 3B, whereas both strains responded equally to Gp2 overexpression, the BW25113:Δ*hfq* strain failed to resume growth even after 10 hours under stasis, whereas the wild-type cells, as expected, resumed growth after 5 hours. Overall, the results suggest that small non-coding regulatory RNAs are involved in the adaptive response to Gp2 mediated perturbation to transcriptional programme because growth resumption fails if small regulatory RNA function becomes curtailed in the absence of Hfq.

**Figure 3.**
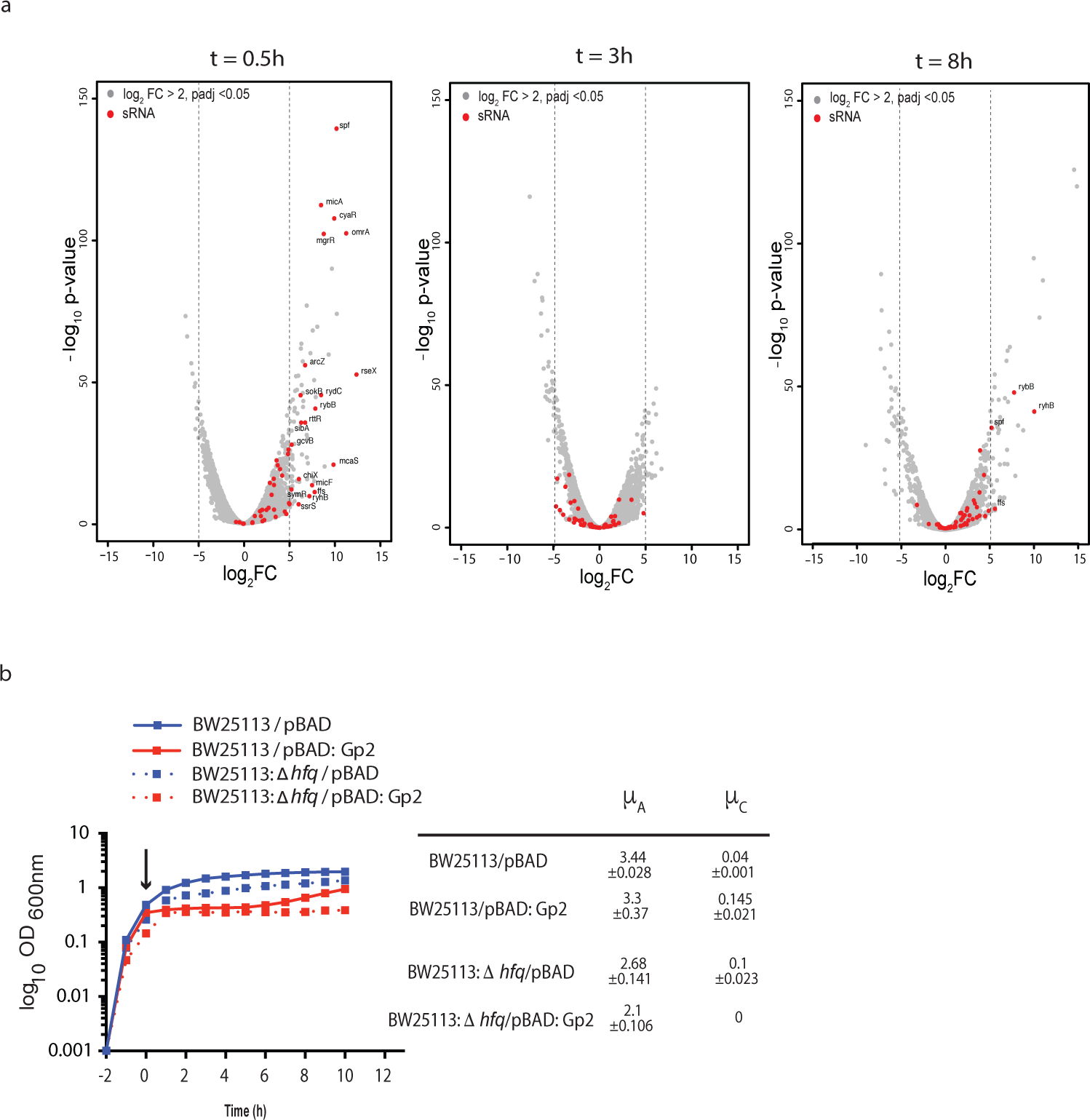
Adaptation to the Gp2 mediated perturbation to the transcriptional programme involves the action of several small non-coding regulatory RNAs. **(A)** The volcano plots, as in Figure 2B, indicating (in red) the expression levels of small non-coding regulatory RNAs. **(B)** Graph showing (log_10_ of the optical density (OD_600_) as function of time (hour)) of a culture of *E. coli* BW25113 and BW25113:Δ*hfq* cells containing pBAD or pBAD:Gp2. The arrow indicates when L-arabinose was added to the culture. The growth rates (μ) of the growing phases of each strain are tabulated on the right of the graph.

### The adaptive response to Gp2 mediated perturbation to the transcriptional programme, at least partly, depends on Eσ^38^

The majority of bacterial adaptive transcriptional responses to stress conditions involve *rpoS*, the gene encoding σ^38^, and Eσ^38^ compared to Eσ^70^ is less sensitive to inhibition by Gp2 (3). Inspired by the observation that induction of Gp2 overexpression in exponentially-growing *E. coli* results in the differential expression of small non-coding regulatory RNA (Figure 3), we sought to understand how expression of small non-coding regulatory RNAs in response to Gp2 overexpression is linked to *rpoS* expression. Interestingly, we note that the expression dynamics of three of the small RNA genes (*arcZ*, *dsrA* and *rprA*) that have a positive influence on *rpoS* expression (18,19) to be substantially up-regulated at t=0.5; however, during stasis, *acrZ*, *dsrA* and *rprA* expression levels substantially drop and increase only moderately during growth recovery (Figure 4A). In contrast, we note that the levels of *oxyS*, which has a negative influence on *rpoS* expression (20), to be moderately up-regulated (by ~2.4 and ~1.5 fold relative to *arcZ* and *dsrA*, respectively) during growth recovery (Figure 4A). Consistent with these observations, analysis of whole-cell extracts prepared from phase B (stasis) and phase C (growth recovery) cells by Western blotting using antibodies against σ^38^, revealed that a marked appearance of σ38 shortly after induction of Gp2 (t=1 in Figure 4B), which rapidly diminished at t=3 (during stasis) and t=10 (upon recovery). Further, consistent with the results in Figure 4B, σ^38^ only co-purifies with Gp2-bound RNAP shortly after Gp2 induction (t=1) and diminishes during stasis (t=3); during growth recovery, at t=10, σ^38^ is not detectably associated with RNAP (Figure 4C). We next analyzed the expression profiles of the σ38 regulon as defined by Weber et al (21) in phases B and C. As shown in Figure 4D, a vast majority of Eσ^38^-dependent genes (58%) were up-regulated shortly after overexpression of Gp2 (t=0.5). However, after 3 hours of stasis (t=3) and upon recovery (t=8) only 37% and 28%, respectively, of Eσ^38^-dependent genes were detected. We next compared the growth characteristics of *E. coli* strain BW25113:Δ*rpoS* with that of the parent strain in response to overexpression of Gp2. Results in Figure 4E show that, although overexpression of Gp2 induces the biphasic growth pattern in wild-type and mutant bacteria, the stasis period in the mutant strain is shortened by ~2 hours compared to that of the wild-type strain; but, both strains resumed growth at the same rate (Figure 4E). Further, we also compared the growth characteristics of *E. coli* strain MG1655:*hfq*(Y25D) with that of the parent strain. The Y25D substitution prevents Hfq from efficiently interacting with *arcZ*, and *dsrA* small RNAs (22) and thus should mimic, to a certain degree, the absence of *rpoS*. The results in Figure 4F clearly indicate that the stasis period in the mutant strain is, as seen with Δ*rpoS* mutant bacteria (Figure 4E), is shortened by ~2 hours compared to that of the wild-type strain (Figure 4F). Overall, the results suggest that *(i)* adaptation to Gp2 induced perturbation to the transcription programme involves, at least partly, stabilization of σ^38^ via the action of small non-coding RNAs *arcZ*, *dsrA* and *rprA*; *(ii)* Eσ^38^ dependent transcription determines the duration of the ‘adaptive’ stasis period, but is not required for establishing stasis and *(iii)* a curtailment of Eσ^38^ activity, possibly but not exclusively *via oxyS*, is required for recovery from stasis.

**Figure 4.**
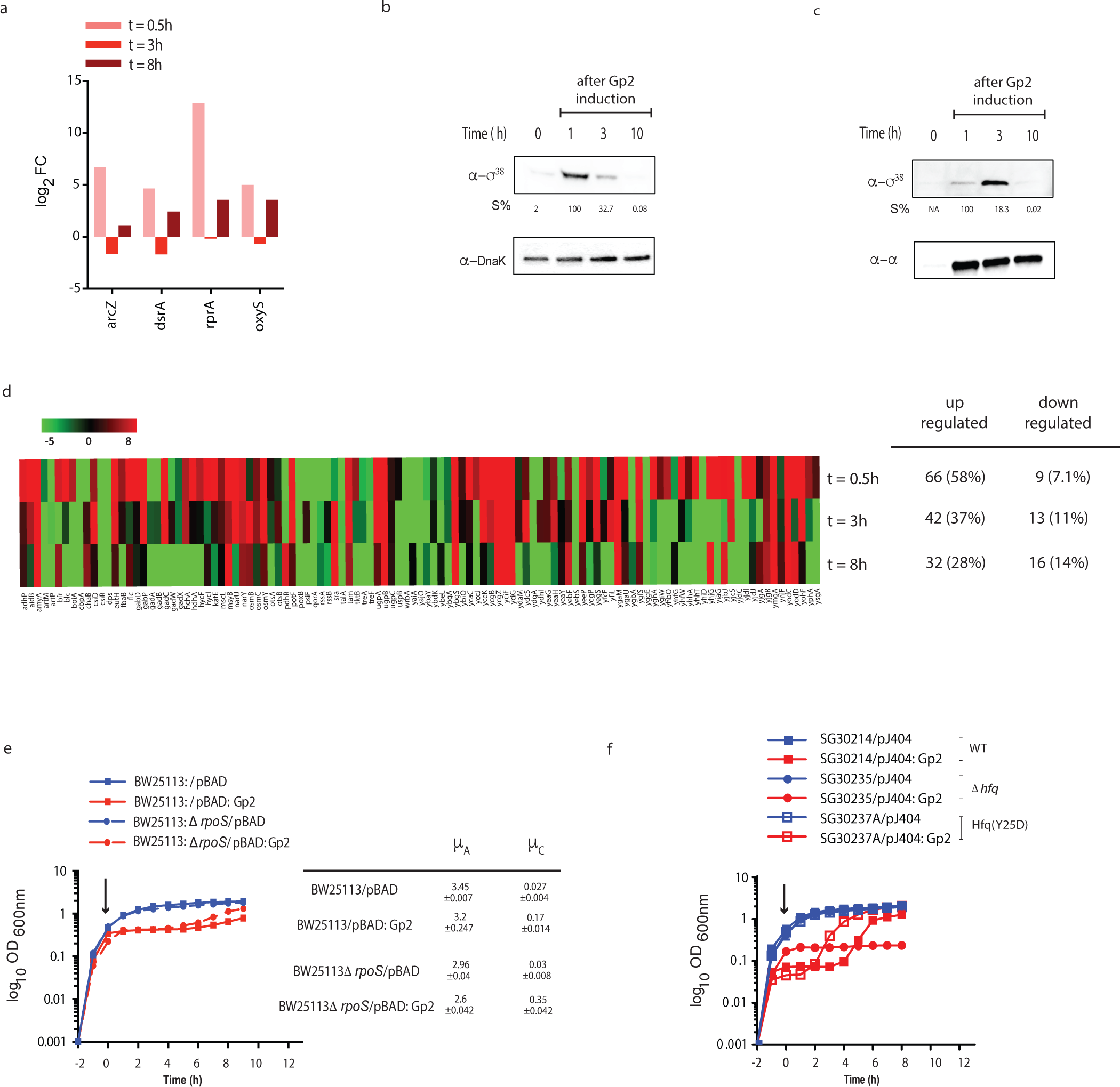
The adaptive response to Gp2 mediated perturbation to the transcription programme, at least partly, depends on Eσ^38^ **(A)** A bar chart showing log_2_ fold change of four small non-coding RNA genes, *arcZ, dsrA, rprA* and *oxyS*, as log_2_ fold change >2 at t=0.5, t=3 (phase B) and t=8 (phase C) relative to gene expression at t=0. **(B)** An image of a Western Blot probed showing σ^38^ levels in whole cell extracts of cells from the indicated time points following induction of Gp2 overexpression; DnaK serves as a loading control. The %S values for σ^38^ bands at each time point is calculated relative to the signal corresponding to the DnaK bands from the same time point and the maximum signal intensity σ^38^ is taken as 100%. **(C)** As in (B) but the image of the Western Blot shows the Eσ^38^ that is bound to Gp2 at the indicated time points (see text for details). **(D)** Heat map showing expression pattern of 113 genes of the σ^38^ regulon (as described in (21)) at t=0.5, t=3 and t=8 relative to t=0. **(E)** Graph showing (log_10_ culture of the optical density (OD_600)_ as function of time (hour)) of a of *E. coli* BW25113 and BW25113:Δ*rpoS* cells containing pBAD or pBAD:Gp2. The arrow indicates when L-arabinose was added to the culture. The growth rates (μ) of the growing phase A and C of each strain are tabulated on the right of the graph. **(F)** As in (E) but the experiment was conducted using *E. coli* SG30214 and mutant derivatives containing pJ404 and pJ404:Gp2 (see Table S1 and see text for details).

### Adaptation to Gp2 mediated perturbation to the transcription programme confers *E. coli* the ability to tolerate diverse antibacterial stresses

Bacteria that display biphasic growth behaviour and recover growth in a heterogeneous manner, for example in response to diauxic nutritional shifts, often display altered susceptibility to antibiotics (23,24). Therefore, since Gp2 mediated perturbation to the transcription programme clearly results in a biphasic growth pattern with a period of no growth (phase B), which precedes a period of slow and heterogeneous growth (phase C), we compared the ability of phase B (t=3 after Gp2 induction) and phase C (t=8 hours after Gp2 induction) cells to survive exposure to ten-fold minimum inhibitory concentration of two different antibiotics (gentamicin and ciprofloxacin) with cells containing the control plasmid from the corresponding time points. As shown in Figure 5A and 5B, surprisingly, after 5 hours of treatment with antibiotics, we observed that phase B exposed to Gp2 displayed ~10 and ~2-fold decreased susceptibility to gentamicin and ciprofloxacin, respectively, compared to the control cells from the corresponding time point. Strikingly, phase C cells that have resumed growth following Gp2 induced stasis, displayed a markedly decreased susceptibility to killing by gentamicin and ciprofloxacin (by ~480 and ~17-fold, respectively) compared to control cells from the corresponding time point. We note that the biphasic nature of the time-kill curve observed with the control cells indicate, unsurprisingly, the presence of persister bacteria in the sample (25). Interestingly, the Gp2 exposed cells display a linear kill kinetic suggesting that Gp2 exposed bacteria are tolerant to antibiotic treatment, as they require more exposure time to be effectively killed than the control cells (25). Additional control experiments confirmed that the Gp2 exposed bacteria (phase B and C) have not acquired genetic changes that allowed them to resist the effect of the antibiotics as the minimum inhibitory concentration (MIC) of each antibiotic required stop them growing remained unchanged compared to control cells (phase A) (Figure 5C). Since the results indicate that Gp2 mediated perturbation to the transcription programme clearly leads to phenotypic changes that enable the cells to tolerate antibiotic exposure, we investigated whether the same was true for other antibacterial stresses, such as exposure to hydrogen peroxide, low pH and high salt. As shown in Figure 5D, 5E and 5F, we observed that phase B cells displayed ~8 and ~5-fold increased ability to survive exposure to hydrogen peroxide and low pH stress, respectively, compared to control cells from the corresponding time point; however, no detectable differences in the ability of phase B cells exposed to osmotic stress was observed (Figure 5E). Similarly, we observed that phase C cells displayed ~33, ~7.5 and ~5-fold increased ability to survive exposure to hydrogen peroxide, low pH and osmotic stress, respectively, compared to control cells from the corresponding time point (Figures 5D, 5E and 5F, respectively). Overall, we conclude that adaptation to Gp2 mediated perturbation to the transcription programme confers *E. coli* the ability to tolerate diverse antibacterial stresses than cell that have not been exposed to Gp2.

**Figure 5.**
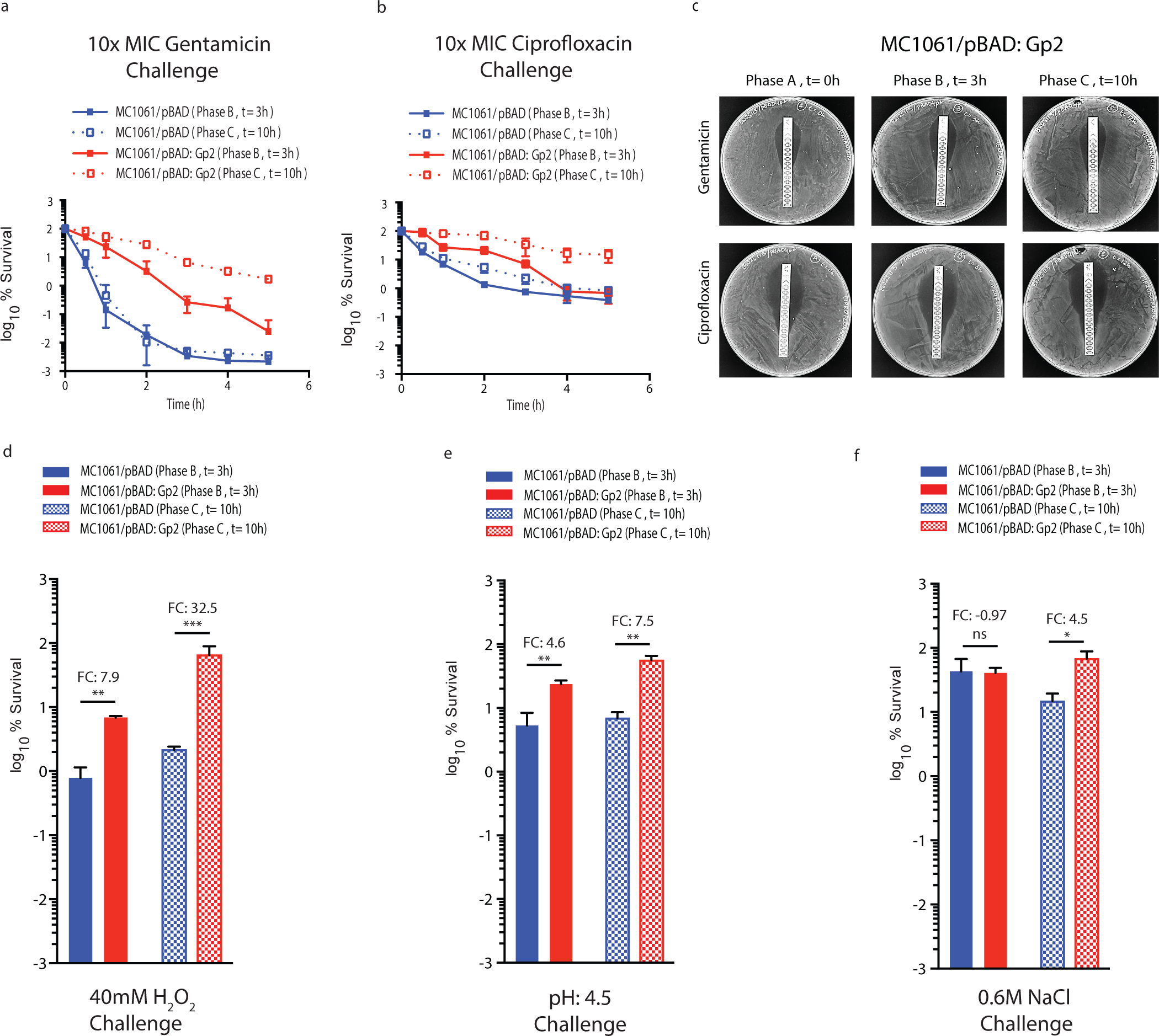
Adaptation to Gp2 mediated perturbation to the transcription programme confers *E. coli* the ability to tolerate diverse antibacterial stresses. **(A)** Graph showing the log10 % survival of *E. coli* MC1061 containing pBAD or pBAD:Gp2 challenged with 10X MIC of gentamicin at t=3 (phase B) and t=10 (phase C) hours after induction of Gp2 overexpression. To calculate log10 % survival the ratio of CFU of untreated cells to and treated cells was obtained and multiplied by 100 for each time point (see materials and methods for details). (B) As in (A) but ciprofloxacin was used. **(C)** *E. coli* MC1061/pBAD:Gp2 cells from t=0 (phase A), t=3 (phase B) and t=10 (phase C) hours after induction of Gp2 overexpression plated on a LB agar plate in the presence of either a gentamicin (top panel) or ciprofloxacin MIC assay strip (see text for details). **(D)-(F)** Bar charts showing log10 % survival of *E. coli* MC1061 cells containing either pBAD or pBAD:Gp2 exposed to 40 mM H_2_O_2_ challenge (D), pH4.5 challenge (E) or 0.6 M NaCl challenge (F) at t=3 (phase B) and t=10 (phase C) relative to untreated cells from the corresponding time point. To calculate log10 % survival the ratio of CFU of untreated cells to and treated cells was obtained and multiplied by 100 for each time point (see materials and methods for details). The error bars on all growth curves represent standard deviation where *n*=3. Statistical significant relationships from One-way ANOVA analysis are denoted (\*\*\*\**P*<0.0001); FC indicates fold-change relative to control (blue bars).

## CONCLUSIONS

The importance of controlling gene expression at the level of transcription in regulating bacterial adaptive responses during early stages of stress exposure is widely established. Transcription factors that detect chemical or physical stress signals within a bacterial cell underpin the regulatory basis of many such adaptive responses. As the only enzyme responsible for cellular transcription, the bacterial RNAP is a nexus for the interaction of transcription factors that direct the reprogramming of cellular transcription to allow bacterial cells to mount the appropriate adaptive responses to changes in growth conditions. As such, the RNAP represents the major mediator of all cellular adaptive processes. The impetus for this study came from our desire to understand how bacteria respond to conditions that specifically compromise RNAP performance. Such a condition can occur during phage infection, action of bacteriocins that target the RNAP, dysregulated transcription factor activity or synthetic modulation transcription networks. The results indicate that a perturbation to the transcriptional programme induced by conditions that compromise RNAP activity can confer bacteria the ability to temporarily and reversibly tolerate exposure to agents that are widely used to control acterial growth. The biphasic nature of bacterial growth and the plasticity of the transcriptome in response to Gp2 indicate that the transcription programme becomes rewired to allow recovery from the condition that has compromised the activity of the RNAP. In the context of the system studied here, it seems that the rewiring of transcription that leads to the recovery of growth, at least partly, but perhaps unsurprisingly, involves the action of small non-coding regulatory RNAs and *rpoS* – the global regulators of bacterial adaptive responses. The mechanistic details underpinning this adaptive response and how the ‘adapted’ cells become refractory to Gp2 (Figure S1) and tolerate the diverse antibacterial stresses (Figure 5) remains unknown, but might not be important in the context of the synthetic system studied here to dysregulate the RNAP. Rewiring of bacterial transcription networks for biosynthetic purposes widely involves the modulating transcription factors and/or promoter function, which often lead to the dysregulation of RNAP performance. Although such engineered bacteria might harbour the intended desirable attribute, importantly, the results of the current study suggest that perturbations to the transcriptional programme caused by dysregulating RNAP function can also confer undesirable traits, such as enhanced tolerance to antibacterial agents. Further, several phages encode small proteins that perturb essential macromolecular processes in bacteria as part of their host acquisition strategy. This study underscores the usefulness of such phage-encoded proteins, like Gp2, as molecular tools to interrogate bacterial physiology and behaviour.

## ACKNOWLEDGEMENTS

We thank members of the S.W. laboratory, Daniel Brown and Martin Buck for constructive comments on the manuscript and Susan Gottesman for strains used in Figure 4F.

## SUPPLEMENTARY FIGURE LEGENDS

**Figure S1.**
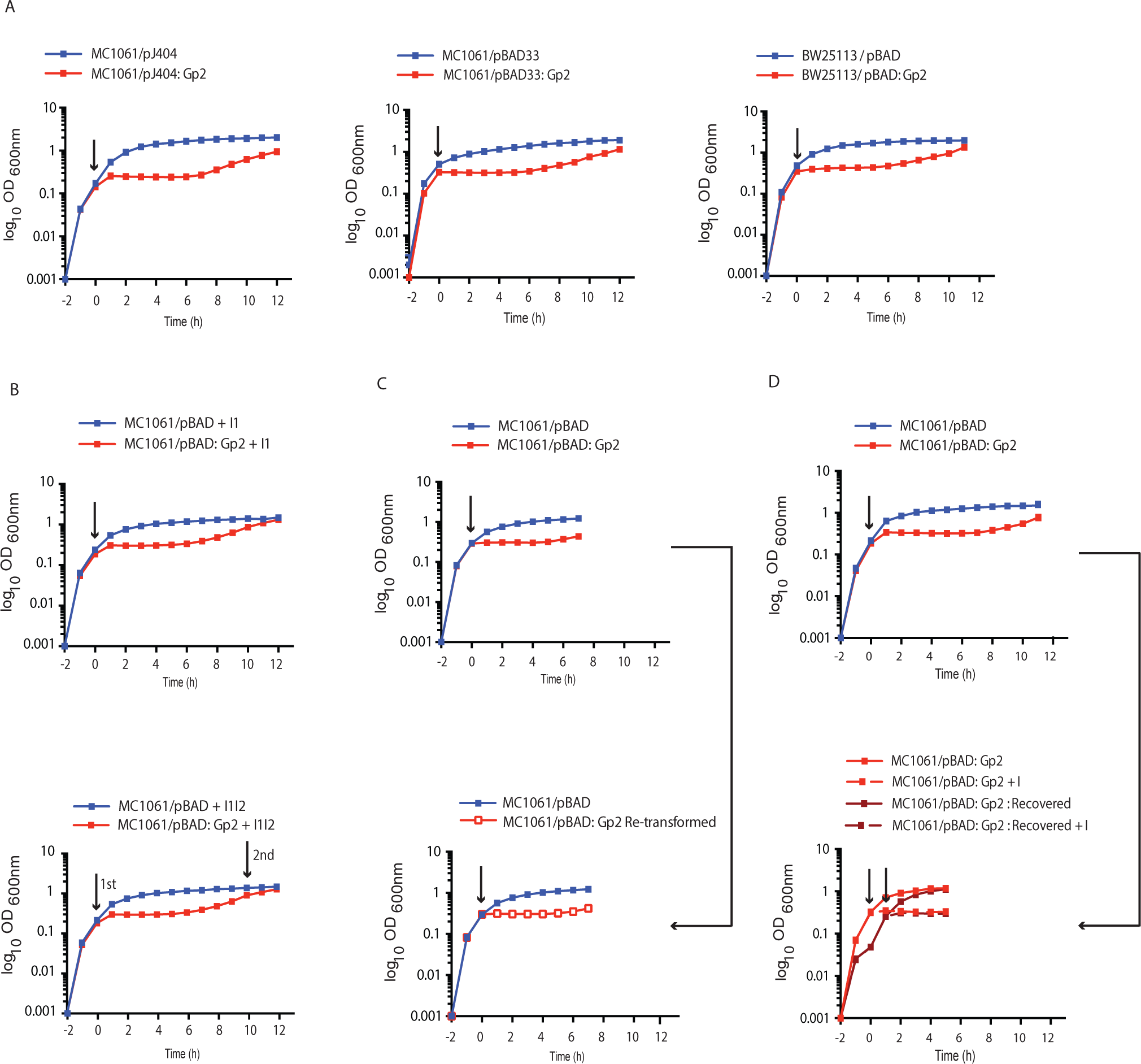
**(A)** Graphs showing (log_10_ of the optical density (OD_600_) as function of time (hour)) of a culture of *E. coli* MC1061 cells containing pJ404 or pJ404:Gp2 (left) and pBAD33 or pBAD33:Gp2 (right). The arrow indicates when L-arabinose was added to the culture. **(B)** As in (A) but the experiment was conducted with *E. coli* BW25113 cells containing pBAD or pBAD33. **(C)** Graphs showing (log_10_ of the optical density (OD_600_) as function of time (hour)) of a culture of *E. coli* MC1061 cells containing pBAD or pBAD:Gp2 where either one (top) or two doses (bottom) of L-arabinose was added at the time points indicted by the arrow (see text for details). The arrows indicate when L-arabinose was added to the culture. **(D)** Graphs showing (log_10_ of the optical density (OD_600_) as function of time (hour)) of a culture of *E. coli* MC1061 cells containing pBAD or pBAD:Gp2 in which both plasmids were isolated at t=8 hours (top graph), transformed into fresh *E. coli* MC1061 cells and grown again as in the top graph in fresh growth medium - shown in the bottom graph (see text for details). The arrows indicate when L-arabinose was added to the culture. **(E)** Graphs showing (log10 of the optical density (OD_600_) as function of time (hour)) of a culture of *E. coli* MC1061 cells containing pBAD or pBAD:Gp2 in which the cells from both cultures were harvested at t=12 (top graph), extensively washed and re-inoculated into fresh growth medium (bottom graph). The arrows indicate when L-arabinose was added to the culture.

**Figure S2.**
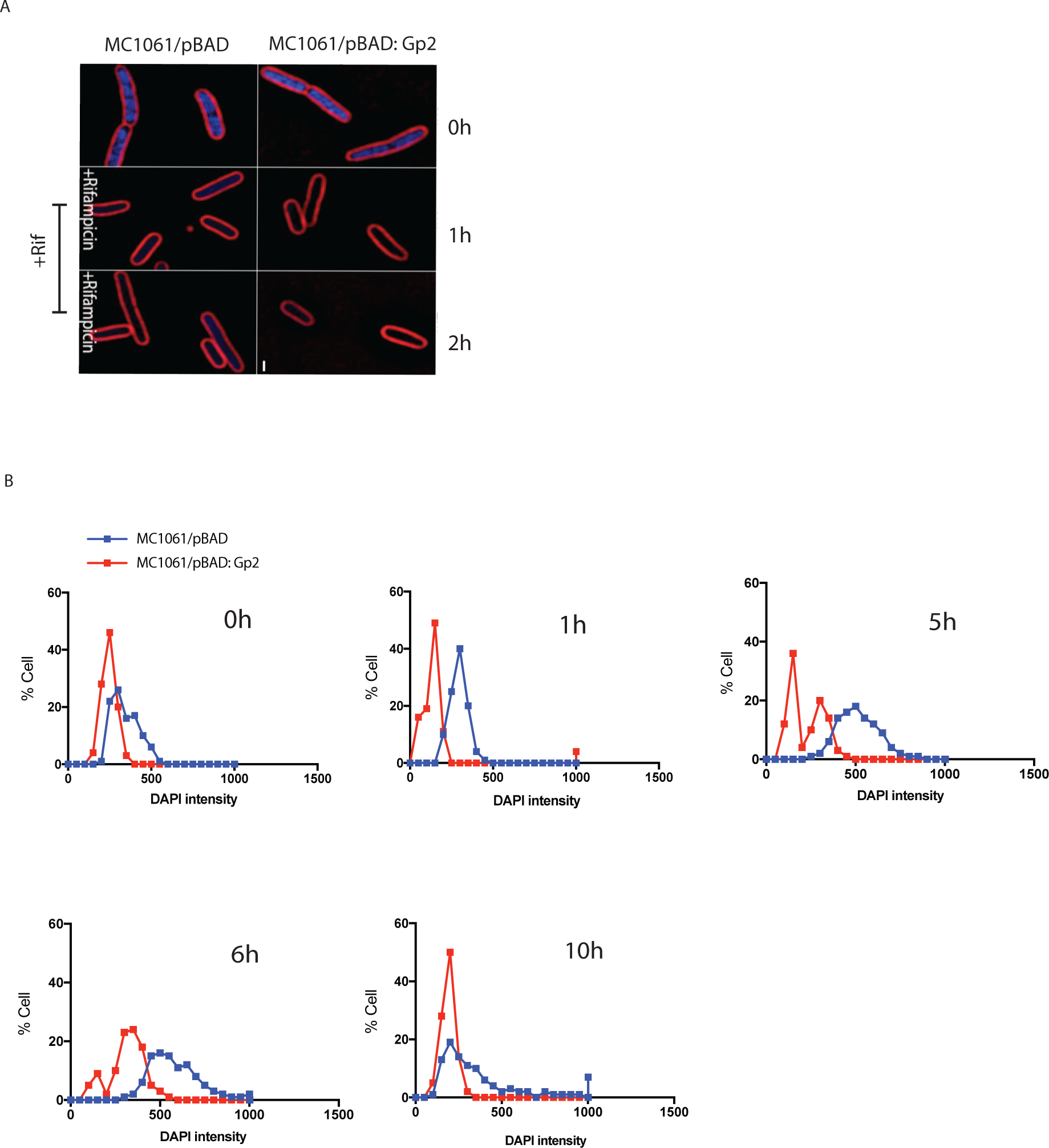
**(A)** Fluorescence microscopy images of cytological profiles of *E. coli* cells containing MC1061/pBAD and MC1061/pBAD:Gp2 before and after treatment with rifamycin. (B) Plot of the intensity of DNA staining (DAPI) in and the recovery of the DNA staining over the course of the experiment shown in Figure 1F (only selected time points are shown).

**Figure S3.**
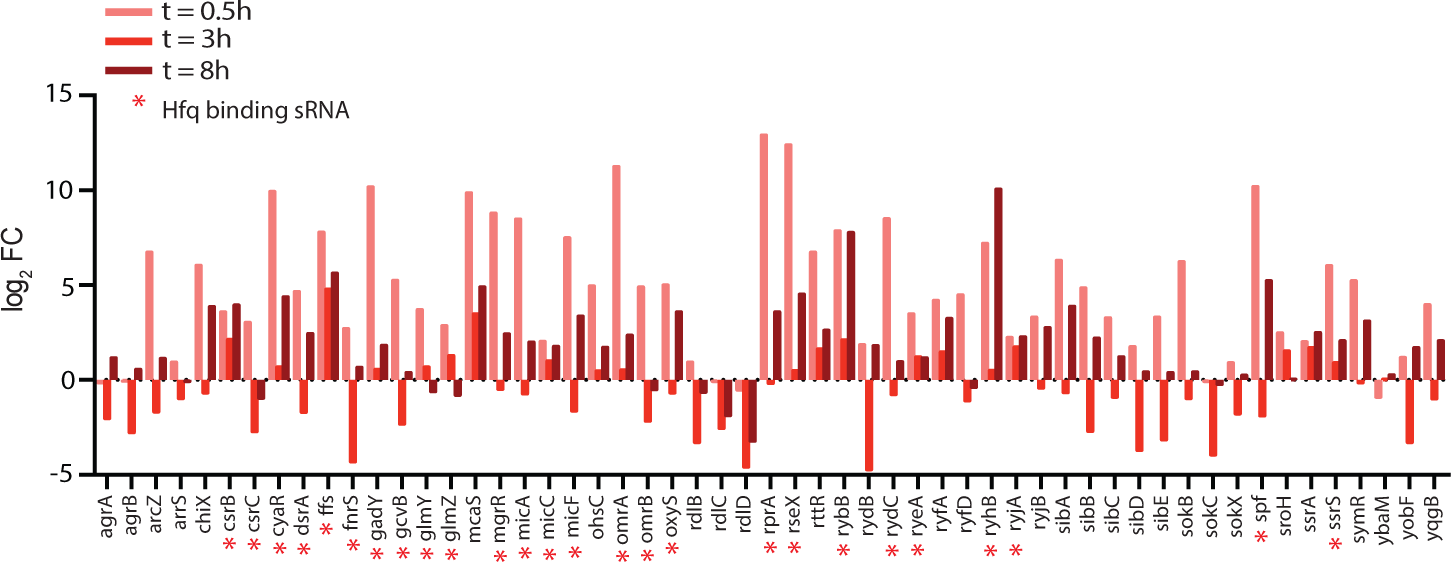
A bar chart showing the log2 fold change of 54 small non-coding RNA genes as log2 fold change > 2 at t=0.5, t=3 (phase B) and t=8 (phase C) relative to gene expression at t=0. The Hfq associated small non-coding RNAs are indicated with red asterisks.

**Figure S4.**
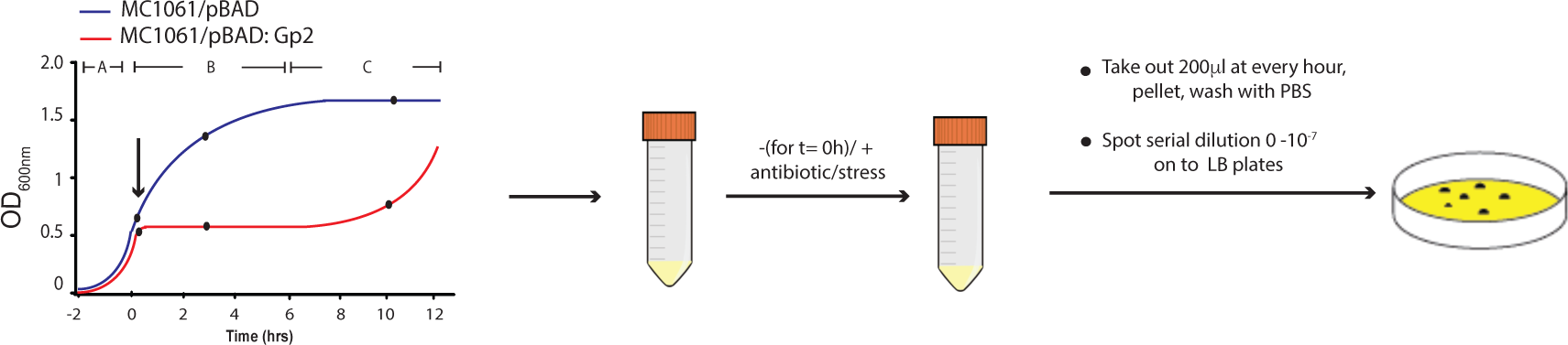
A schematic showing the experiment done for determining the survival of *E. coli* cells containing MC1061/pBAD and MC1061/pBAD:Gp2 at t=3 (phase B) and t=8 (phase C) following antibiotic or stress exposure (see text and Materials and Methods for details)..

**Supplementary Table 1.**
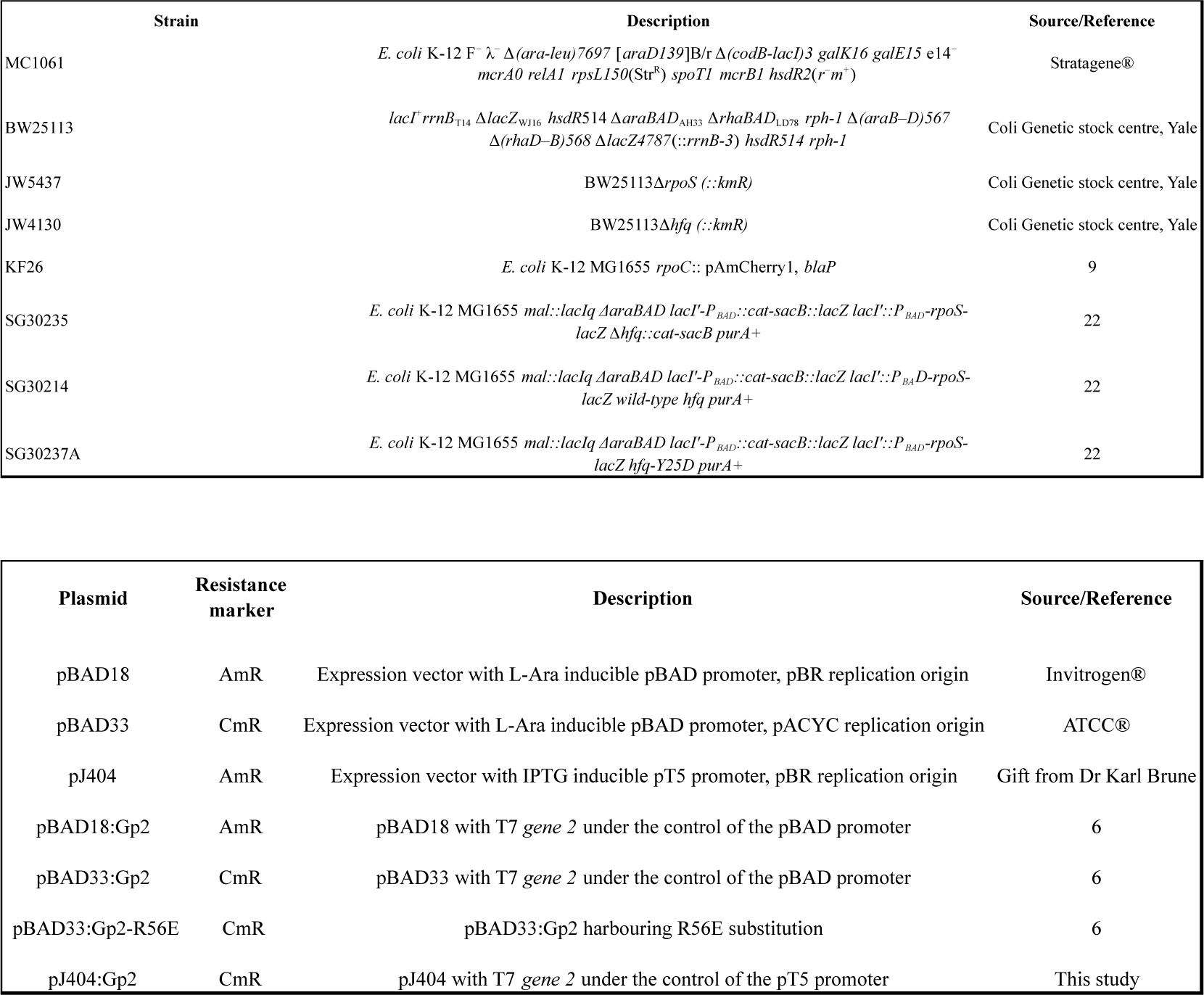

